# Generated randomly and selected functionally? The nature of enterovirus recombination

**DOI:** 10.1101/2020.09.29.319285

**Authors:** Fadi G. Alnaji, Kirsten Bentley, Ashley Pearson, Andrew Woodman, Jonathan Moore, Helen Fox, Andrew MacAdam, David J. Evans

## Abstract

Genetic recombination in RNA viruses is an important evolutionary mechanism. It contributes to population diversity, host/tissue adaptation and compromises vaccine efficacy. Both the molecular mechanism and initial products of recombination are relatively poorly understood. We used an established poliovirus-based *in vitro* recombination assay to investigate the roles of sequence identity and RNA structure, both implicated or inferred from analysis of circulating recombinant viruses, in the process. In addition, we used next generation sequencing to investigate the early products of recombination after cellular co-infection with different poliovirus serotypes. In independent studies we find no evidence for a role for RNA identity or structure in determining recombination junctions location. Instead, genome function and fitness are of greater importance in determining the identity of recombinant progeny. These studies provide further insights into this important evolutionary mechanism and emphasise the critical nature of the selection process on a mixed virus population.

## Introduction

The rapid evolution of RNA viruses is attributable to both the error prone nature of viral RNA-dependent RNA polymerases (RdRp) and the extensive exchange of genetic information achieved through the processes of recombination and – in segmented viruses – reassortment. Together, these are important drivers of virus evolution, often linked to the emergence of novel pathogens and disease outbreaks [1-4]. Recombination predominates in the single stranded positive-sense (mRNA-sense) RNA viruses and is proposed to occur via distinct replicative or non-replicative mechanisms. In the latter the formation of a full-length genome likely involves processing and ligation by cellular enzymes, and the detailed mechanism and biological significance remain unclear [5, 6]. In contrast, replicative recombination between two compatible virus genomes co-infecting the same cell is increasingly well-studied [7-10].

The importance of recombination in nature is typified by the frequent isolation of recombinant forms of members of the *enterovirus* genus of the family *Picornaviridae*. Poliovirus and other picornaviruses have been extensively used to study recombination and the underlying mechanisms involved [8, 11]. These include the initial demonstration of the copy-choice strand-transfer event and numerous, but often conflicting studies interpreting the influence of RNA sequence and structure on the generation of viable recombinants [10, 12-17]. To facilitate the analysis of early events in recombination we previously established an *in vitro* assay (the CRE-REP assay) that solely yields recombinant viruses [18].

To directly address the potential influence of RNA structure and sequence identity on the strand-transfer event in recombination we have exploited the CRE-REP assay and modified the input genomes. By independently analysing genomes containing extensive regions of sequence identity, or templates modified to contain regions of high or low levels of RNA structure, we were unable to demonstrate any significant influence on the proportion of recombination events occurring in these modified regions. We extended these studies to investigate the early RNA products arising from a natural co-infection of cells with two serotypes of poliovirus using deep sequencing. In contrast to the generally clustered range of recombinants found using the CRE-REP assay [18] these were distributed throughout the genomic region analysed. Based on our analyses we propose that neither sequence identity nor RNA structure are important primary determinants in recombination.

## Results

### Influence of RNA structure and sequence on CRE-REP recombination

The CRE-REP assay [18] utilises a poliovirus type 1 (PV1) luciferase-encoding subgenomic replicon as the polymerase donor and a poliovirus type 3 (PV3) genome bearing a lethal mutation in the *cis*-acting replication element (CRE) located within the 2C-coding region [19, 20] as the acceptor (**Fig 1A**). Co-transfection of *in vitro*-generated RNA from both templates only yields viable recombinants if a suitable strand-transfer event 5′ to the functional CRE element in the donor and 3′ to the P1 capsid-coding region in the acceptor template occurs. We previously demonstrated that viable recombinants had junctions that predominantly clustered in one of two regions, spanning either the VP1/2A or 2A/2B polyprotein cleavage boundaries (Cluster 1 and Cluster 2 respectively, hereafter known as C1 and C2). We reasoned that by modifying the sequence identity or RNA structure within the C2 region of the donor or acceptor templates, we could investigate the local influences of sequence or RNA structure on recombination using the unmodified C1 region as an internal control. For example, a positive influence on recombination would yield a higher ratio of C2:C1 recombinants than observed in a control assay with unmodified templates.

**Fig 1.**
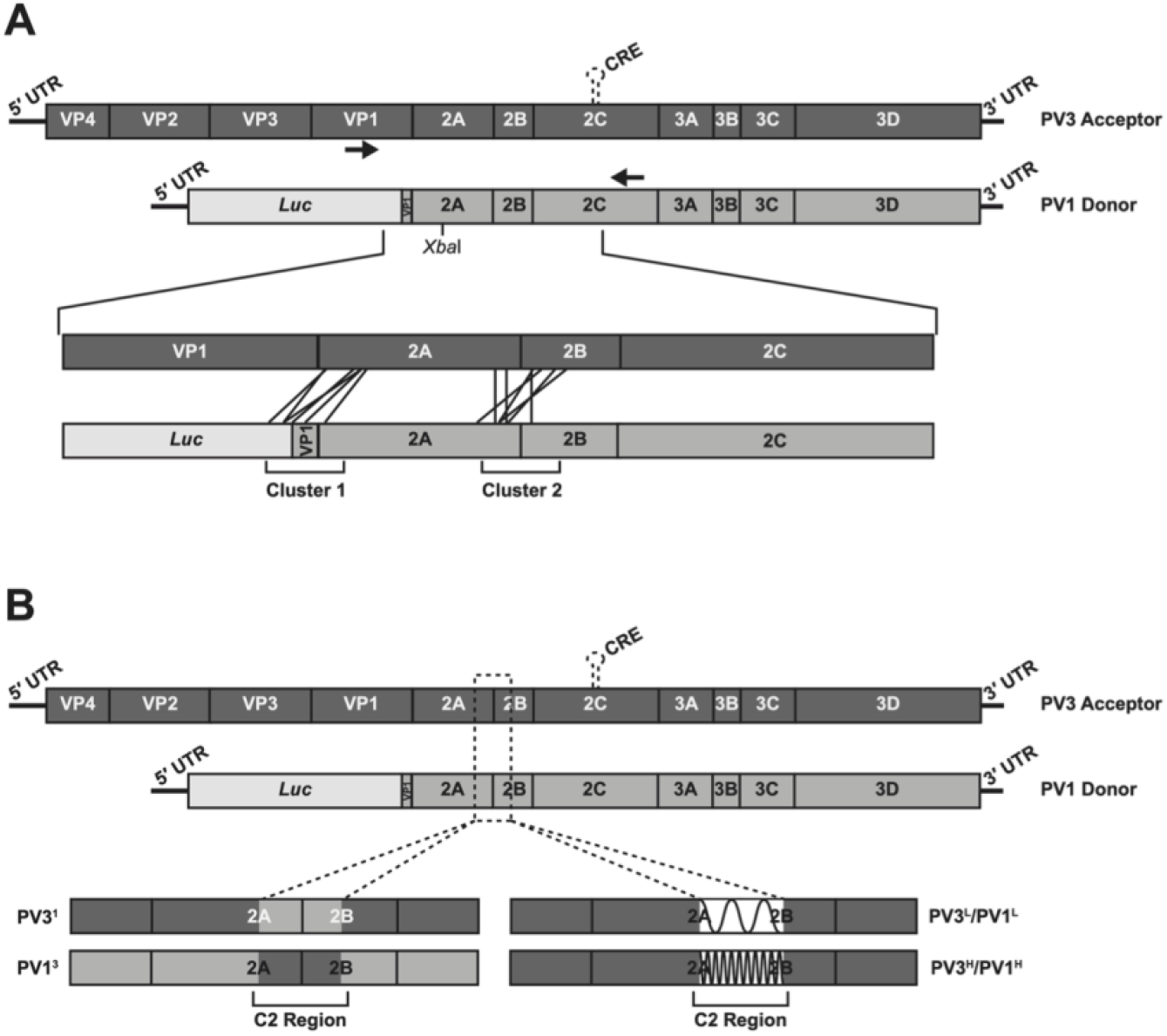
Schematic representation of CRE-REP assays. **(A)** Intertypic CRE-REP assay showing the genomes of the PV3 acceptor and PV1 donor RNAs. Black arrows represent position of primers used to amplify recombinant genomes. The lower expanded image illustrates the recombination window and highlights examples of recombination events in Cluster 1 and Cluster 2. The position of a unique *Xba*I site used for screening is shown in the donor template. **(B)** Modified CRE-REP assays highlighting the region of modification within Cluster 2 and the resulting acceptor and donor templates with altered sequence identity or RNA secondary structure.

We exploited the redundancy of the amino acid triplet code to design 450 nt sequences, centered on the 2A/2B cleavage site, with high or low levels of RNA structure (see Materials and Methods). All designed sequences exhibited the same level of sequence identity when compared to their parental sequence (85.5%) and retained the same level of sequence identity as present between the parental PV1 and PV3 CRE-REP partners (77.6% +/-1%; Table S1). In each case the encoded proteins were identical to the respective native PV1 donor or PV3 acceptor templates. The resulting modified RNA templates used in CRE-REP assays were: (1) the PV3-derived acceptor modified PV3^L^ and PV3^H^ (where L and H refer to low and high RNA structure respectively) and (2) the PV1-derived donor modified PV1^L^ and PV1^H^ (**Fig 1B and S1**).

Similarly, we engineered donor and acceptor templates such that the same 450 nt region was ∼98% identical. This was achieved by directly exchanging the PV1 sequence with that of PV3, or vice versa, with the exception of retention of the seven amino acids that differ between PV1 and PV3 in this region (see Materials and Methods) to avoid disruption of any *in cis* interactions involving these residues. The resulting modified RNA templates for use in CRE-REP assays were PV3^1^ (a PV3-derived acceptor containing a stretch of PV1 sequence), and PV1^3^ (a PV1-derived donor containing a stretch of PV3 sequence; **Fig 1B and S1**). We confirmed that the sequences introduced to the CRE-REP donor and acceptor templates had no significant effect on virus replication (**Fig S2**) and therefore should not influence the underlying replicative recombination process.

To generate recombinant virus populations, we conducted CRE-REP assays in parallel. For each assay one unmodified template (either donor or acceptor) was paired with one modified donor or acceptor template. Assays are referred to by the name of the modified template of the RNA pair. Donor and acceptor RNAs were co-transfected into murine L929 cells (permissive but not susceptible to poliovirus) in equimolar amounts and the virus-containing supernatant harvested 30 hour post-transfection. In addition, we transfected each donor and acceptor RNA individually into L929 cells and confirmed that, as expected, none were alone able to generate infectious virus (data not shown). A total of three independent co-transfections were carried out for each assay and the yield of recombinant virus determined by plaque assay (**Fig S3**). The unmodified CRE-REP control assay (WT) generated an average of 2.1 × 10^3^ pfu/ml of recombinant viruses. The total yield of recombinants from CRE-REP assays with template modifications did not significantly differ from the WT control assay, demonstrating no obvious advantage or disadvantage in recombinant generation resulting from the modifications.

### Analysis of recombination junctions in the modified CRE-REP assay

The overall yield of recombinants (**Fig S3**) includes all those that map in C1 and C2 combined. To determine whether there were any changes in the ratio of recombination events occurring within the C1 and C2 regions as a result of the introduced modifications we sequenced a representative sample of recovered viruses. The supernatants from the triplicate transfections of each assay were pooled to generate a diverse virus population and minimise founder-effects. A total of 420 recombinants (60 from each CRE-REP assay) were isolated by limit dilution. Following viral RNA extraction and reverse transcription, the entire recombination region was amplified by PCR and sequenced to determine the location and features of each recombination junction (**Fig 2A**).

**Fig 2.**
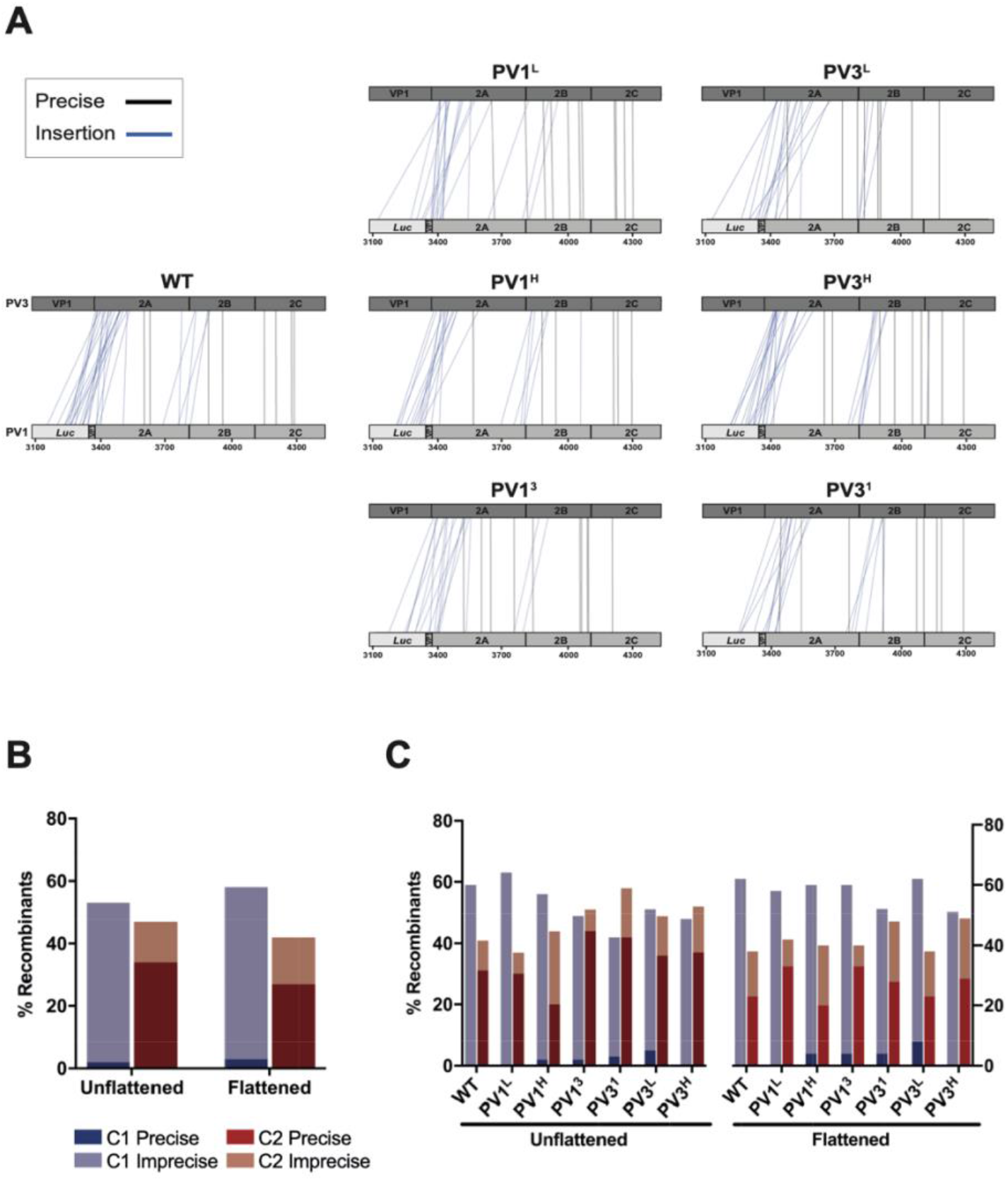
Analysis of recombination junctions from CRE-REP assay. **(A)** The location of each recombination junction was mapped respective to each parental genome. Each line represents a unique recombinant within the population of precise (black) or imprecise-insertion (blue) recombinants (see Nomenclature, materials and method). **(B)** Junctions were defined as precise or imprecise and graphed according to location in Cluster 1 or Cluster 2. Unflattened data was compared to flattened data, using Fishers Exact Test and showed no significant difference in the distribution of recombinants. **(C)** Junctions were defined as for (B) and graphed by individual CRE-REP assay. Each modified assay was compared to wild type using Fishers Exact Test and showed no significant difference in the distribution of recombinants, either between assays or unflattened and flattened data sets.

After sequencing, 119 samples were discarded due to ambiguities in the sequence that indicated the virus population was not clonal. Approximately one third of the remaining recombinants were present more than once in the dataset. These identical sequences could either arise independently due to unique recombination events or may reflect early recombination products that have undergone additional rounds of replication (founder-effects), so increasing their proportion in the mixed virus progeny. Since the experimental design could not discriminate between these, we considered both the data in its entirety (unflattened) and after discarding identical sequences (flattened).

In the flattened data (n=205) 58% of all junctions mapped to the C1 region while 42% mapped to the C2 region. No statistical difference (Fishers Exact Test) was observed when compared to the unflattened data (n=301), where 53% mapped to C1 and 47% to C2 (**Fig 2B**). Further analysis of the data demonstrated that there were also no statistically significant (Fishers Exact Test) favoured or disfavoured modified assay pairings that altered the distribution of junctions between the C1 and C2 regions when compared with the unmodified WT templates (**Fig 2C**).

These results suggested that neither local sequence identity, nor the gross level of RNA structure, play a major role in influencing the location of recombination junctions within viable recombinants that were selected, packaged and subsequently propagated.

### Isolation of recombinants following virus co-infection

If, as we suggest, RNA sequence and structure do not influence the process of recombination we would expect the junctions to have been functionally selected i.e. on viability or relative fitness, from a much more diverse spectrum of crossover events. Such a diverse population potentially includes both in-frame insertions and deletions and out-of-frame junctions. Of these, insertions have been previously reported [18] but deletions or out-of-frame recombinants are highly unlikely to be present amongst viable recombinant progeny. To investigate recombinant genome diversity further we developed an assay based upon the next generation sequencing (NGS) analysis of a mixed virus population arising after co-infection with poliovirus serotypes 1 and 3.

HeLa cells were co-infected at an MOI of 10 and intracellular viral RNA was harvested 5 hours post-infection (hpi), a duration previously shown to allow the production and detection of intracellular recombinants [19], whilst minimising the selection and analysis of genomes that have been packaged or initiated further rounds of infection. Purified RNA was reverse transcribed, and PCR amplified across the region spanning Clusters 1 and 2, before analysis by gel electrophoresis (**Fig 3A**). A distinct product of ∼1.3 kb, representing the expected size of a precise recombination product was observed, together with both larger (up to ∼3 kb) and smaller (∼0.45 kb) products. In addition, there was extensive diffuse background staining between these products suggestive of a range of different sized PCR products (**Fig 3A)**. Preliminary analysis of 7 randomly-selected cloned PCR products by Sanger sequencing (**Table S2**) showed a diversity of recombination junctions and encouraged us to study the population of molecules using NGS.

**Fig 3.**
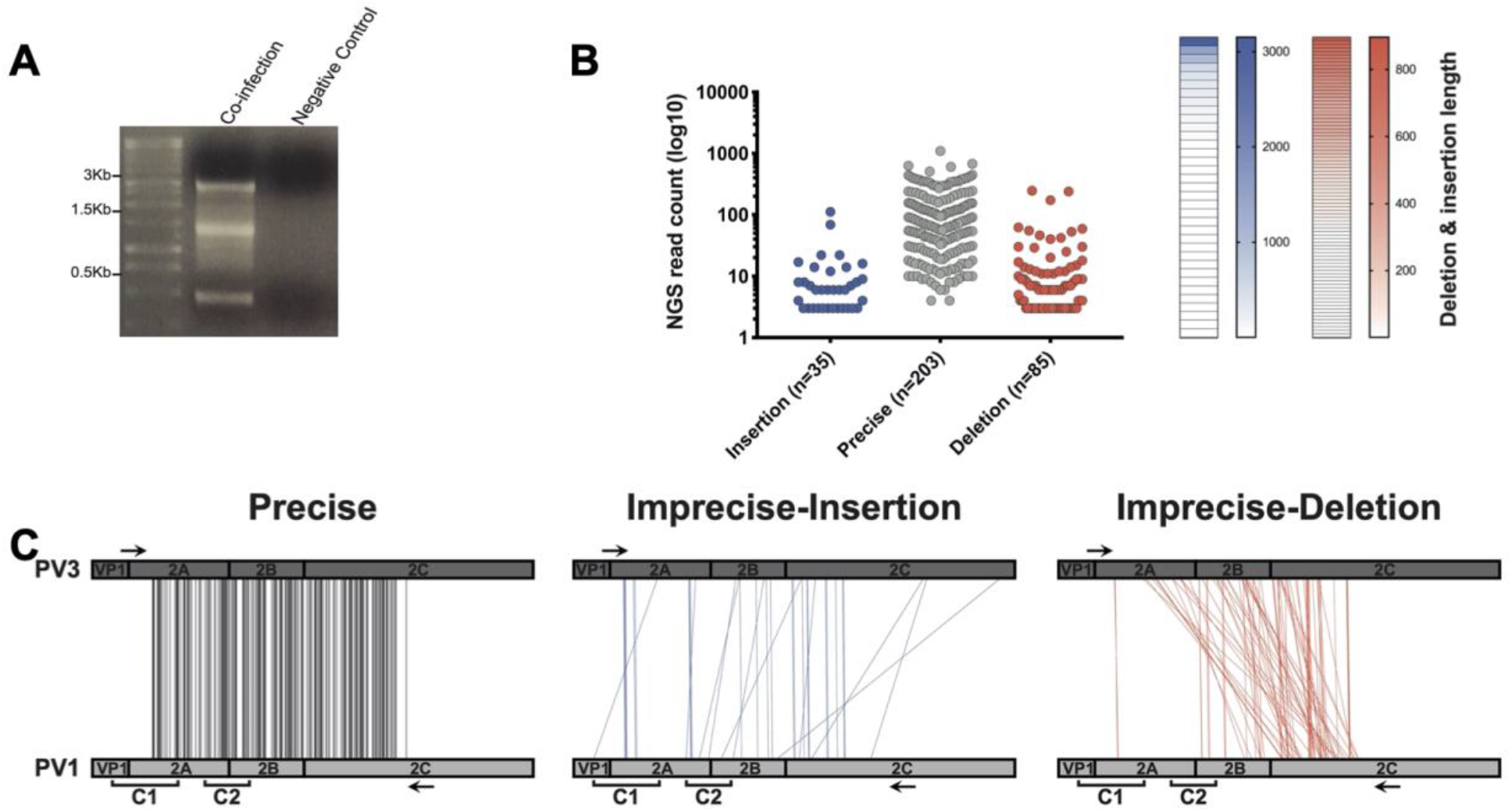
Isolation and characterization of recombinants generated from virus co-infection. **(A)** PCR amplification of the region between nucleotides 3235-4548 followed by agarose gel screening. The negative control is an equimolar mixture of *in vitro* transcribed full-length PV1 and PV3 RNA **(B)** The number of precise and imprecise (both insertion and deletion) recombinants was plotted against the number of NGS reads (left panel); each dot represents a unique recombinant. On the right panel, heatmaps show the distribution of sequence lengths of insertion (blue) and deletion (red) within the detected imprecise recombinants. Each cell represents a unique recombinant. **(C)** Parallel coordinates visualization of recombinants from co-infection studies. The location of each recombination junction was mapped respective to each parental genome. Each line represents a unique recombinant within the population of precise (black), imprecise-insertion (blue) and imprecise-deletion (red) recombinants.

### The complexity of recombination events revealed by NGS sequence analysis

After the recombination-specific amplification of the viral RNA 5 hpi, the amplicons were fragmented before being sequenced on a MiSeq Illumina platform, generating ∼1.5 million reads of 250 nts length. We developed a pipeline based upon the aligner Bowtie2 [20] and the Viral Recombination Mapper (ViReMa) [21] to analyse recombination junctions, and used it as previously described [22]. Briefly, the entire dataset was aligned to WT PV3 and PV1 reference sequences using Bowtie2 and those that matched perfectly i.e. outwith the recombination junction, were eliminated. Remaining reads were analysed using ViReMa to identify the recombination junction (**Table 1**).

**Table 1.**

|  | Negative Control | Co-infection | Technical Replicate |
| --- | --- | --- | --- |
| Total reads | 797386 | 1488344 | 1023536 |
| Aligned to PV1 | 11754 | 279958 | 46028 |
| Aligned to PV3 | 525214 | 1111733 | 909263 |
| Not Mapped* | 260418 | 69292 | 61749 |
| Recombination reads<br>(by ViReMa) | 0 | 27361 | 6496 |

\* Reads derived from cellular genome or tagged 'unknown' for a variety of reasons, including high levels of sequence variation and premature read truncation

We identified ∼27 × 10^3^ recombinant junction sequences (**Table 1**) in the dataset, defining 323 unique junctions, between PV3 and PV1. There were no recombinant junctions in the control dataset. Identified recombinant junctions were initially grouped into those with no sequence duplication or deletion (so called precise junctions [18], n=203; see nomenclature in Materials and Methods), those with sequence insertions (imprecise insertions, n=35), or those with sequence deletions (imprecise deletions, n=85) (**Fig 3B**). The two groups of imprecise junctions exhibited a range of different sized insertions (1 – 3150 nt) or deletions (1 – 896 nt; **Fig 3B**). All 323 recombinants were mapped using parallel coordinate diagrams, which clearly showed that recombination junctions, whether precise or imprecise, were distributed throughout the analysis region (**Fig 3C**), with no evidence for the clustering around the encoded polyprotein cleavage sequences seen in previous CRE-REP analysis [18].

To validate these observations, we performed a technical replicate by independently extracting total cell RNA from the original co-infection sample (**Fig S4, Table 1**). Despite fewer NGS reads overall, and a concomitant reduction in ViReMa-detected junctions, we were still able to identify hundreds of recombinants between PV3 and PV1. Correlation analysis of the NGS reads for each junction in the two replicates, normalised to the total reads per junction, demonstrated the method produced statistically reproducible results (Spearman R=0.88). All subsequent analysis used the first sequenced population containing 323 unique junctions.

### Analysis of recombination junctions following NGS

The NGS analysis of junction sequences provides an independent approach to determine the randomness, or otherwise, of the recombination process. To formally test whether the observed crossover junctions between PV3 and PV1 were randomly distributed across the targeted region, we compared the precise junction locations with a modelled dataset containing randomly generated precise junctions. The comparison used a sliding cumulative score across the window of recombination to determine the occurrence of recombination events at each nucleotide position (**Fig 4A**). We observed no significant difference between our NGS dataset and the randomly generated dataset (P=0.06; Mann–Whitney U test), supporting our contention that precise recombination junctions are randomly located throughout the targeted region.

**Fig 4.**
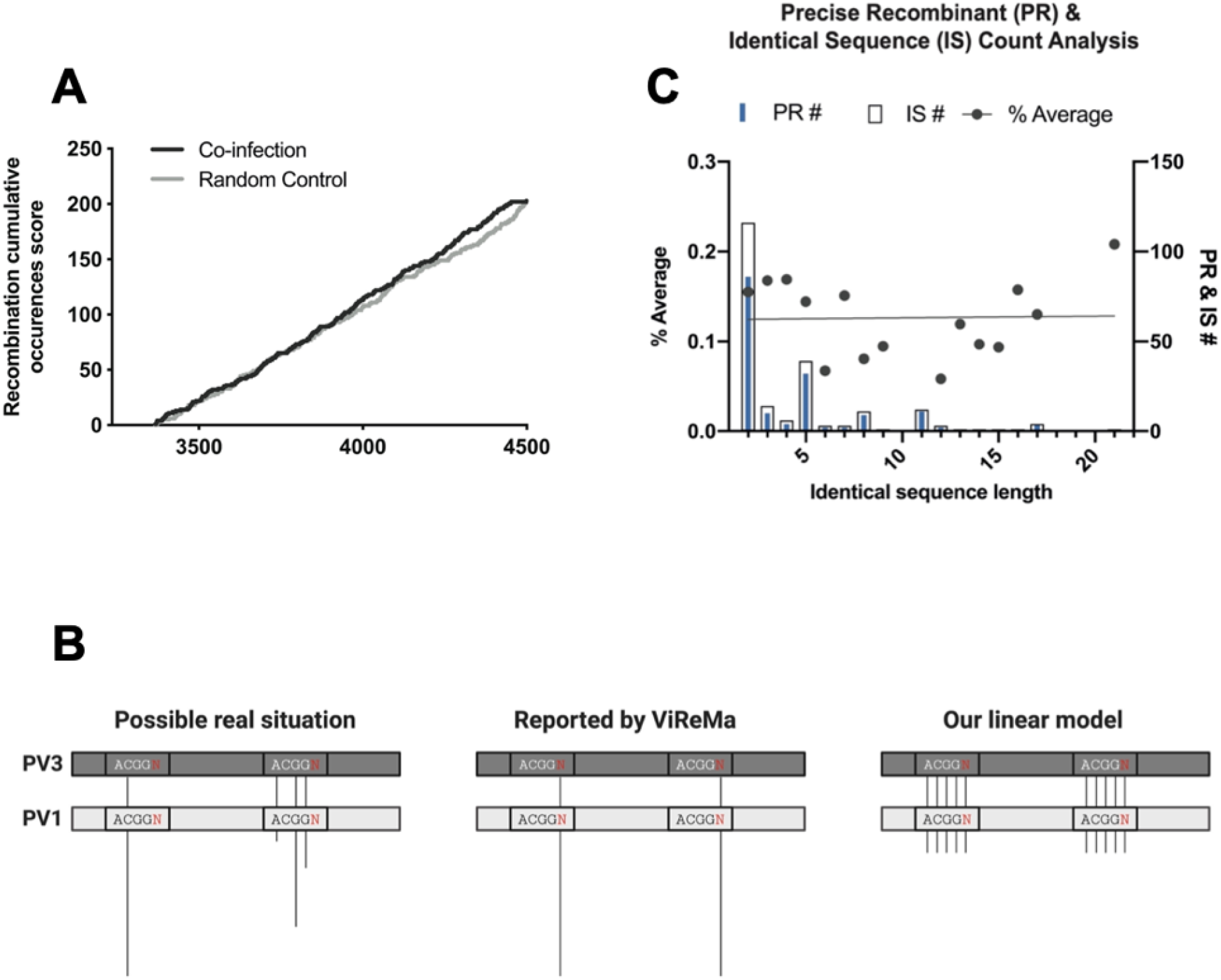
Analysis of recombination junction sequences generated from virus co-infection. **(A)** The occurrence of recombination on the genome was given a score of 1 while no occurrence is 0. The number of recombination occurrences was calculated in a cumulative way i.e. the occurrence of recombination at each position is added to the occurrence at the previous position. This was applied to two populations; the precise recombinants from the virus co-infection sample and a random recombination model generated by Excel. The cumulative recombinant count (y-axis) was plotted against the locations on the PV1 (donor) amplicon (x-axis). **(B)** An Identical sequence length (IS) of 4-nt that hypothetically occurred twice in the genome was used as an example. The rectangles correspond to a part of the targeted region of PV1 and PV3, the lines represent recombinants and their lengths denote the NGS read. i.e. the longer the line the more NGS reads. The identical sequence is placed within a smaller rectangle and the red Ns represent a mismatch of any other nucleotides. Looking at a possible real situation in the left panel, the Precise Recombinants (PRs) occurred at different sites within the identical sequence with different reads. ViReMa will always report 1 PR per IS; something like the middle panel. In our linear model we convert ViReMa model into the right panel, where junction counts equal the number of sequence identity length + the mismatch, and NGS reads are equal for all junctions. **(C)** Individual counts of identical sequences (IS) of different lengths were summed within the amplified region and plotted along with the count of precise junctions that mapped to each different length of sequence identity. For each of the latter the average number of junctions per nucleotide was calculated assuming recombination was equally likely to occur at each position within identical donor and template sequences. The linear regression was calculated to compare the slopes between the fitted line and the perfect line.

As short regions of localized sequence identity have previously been suggested to influence the recombination process [10, 12, 23, 24], we next investigated a role for such sequences in shaping our NGS dataset. Within the targeted region there are 219 short regions of sequence identity between 2 and 32 nt in length. ViReMa maps junctions to the first divergent nucleotide 3′ to a region of identity [21], which makes it impossible to determine the exact nucleotide – within the identical sequence – at which strand transfer occurred. Because of this, we made the assumption that recombination was equally likely to occur between any nucleotide within a region of sequence identity and its adjacent 3′ nucleotide (**Fig 4B**). We normalized the recombinants per nucleotide on this assumption; for example, if a precise junction with 100 NGS reads mapped 3′ to a 4 nt region of aligned PV3 and PV1 sequences, this equated to five precise junctions each represented by 20 reads (**Fig 4B**).

Notably there were no recombinants that mapped to the most extensive region (32 nt.) of sequence identity between the parental genomes. Of the remaining 218 regions (2 - 21 nt.), with reads normalised as described, there was no significant difference (P=0.93) in linear regression analysis of the fitted line for the observed data or the perfect line (**Fig 4C**). This suggests, for precise junctions at least and at this time point of the co-infection of poliovirus serotypes 1 & 3, that recombination junctions are distributed independently of sequence identity.

We additionally failed to detect enrichment of short regions of sequence identity at imprecise junctions (**Fig 5A)**. Correlation analysis between the experimental data and an *in silico* library containing every possible imprecise recombinant within the targeted region, showed a significant positive correlation (R =0.8, p<0.01). This suggests that imprecise recombination tracks closely with the random model and indicates no role for sequence identity in their generation. Finally, we mapped all the imprecise recombinants to their genome positions and we could not detect any correlation between the NGS read frequency and the length of insertion/deletion sequences, reading frame, or location on the genome (**Fig 5B**).

**Fig 5.**
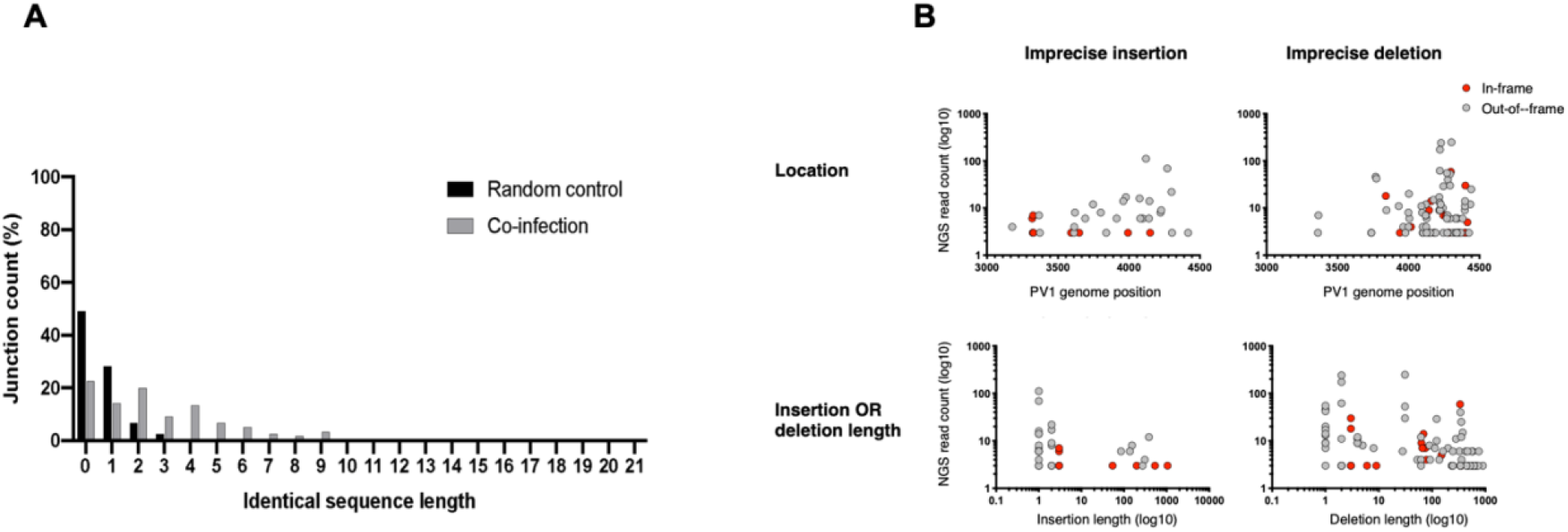
The influence of identical sequence length and reading frame on imprecise recombination. (A) The imprecise recombinants were counted at each identical length (x-axis) and the percentage proportion related to the total imprecise junctions was calculated (y-axis). The flattened data of the imprecise recombinants generated from the co-infection experiments (grey) was plotted against a random control (black), where every possible imprecise junction was created and classified using a custom perl script. (B) In the upper panel, the location of the detected imprecise recombinants on the PV1 genome was plotted against the NGS read count. In the lower panel, the size of either the insertions or deletions of the detected recombinants were plotted against NGS read count. Each dot represents a unique recombinant.

While the above analysis strongly suggested that recombination junctions were randomly distributed, it remained a possibility that localized regions of RNA structure may influence junction location. To investigate these possibilities, we applied mean folding energy difference (MFED) analysis [25] to all detected recombinants of all types. In comparing the MFED values from either the positive-or negative-strand to the number of recombination junctions occurring within the same sliding window, we were unable to detect any correlation between the presence, or absence, of predicted RNA structure and the location of recombination junctions (**Fig 6**). This was in agreement with the CRE-REP assay analysis (**Fig 2**) and further supports our view that RNA structure does not influence the location of recombination junctions.

**Fig 6.**
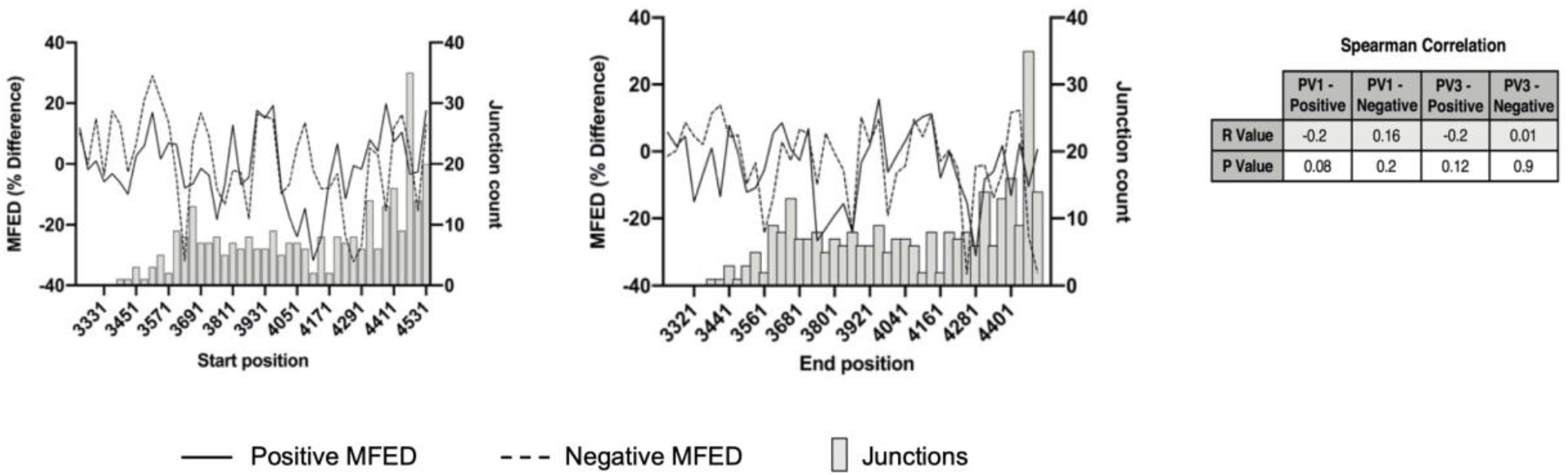
The influence of RNA structure on recombination. The MFED was calculated for a 100 nt sliding window (at 30 nt increments) for both PV1 and PV3 positive and negative strands (dashed and solid lines respectively) and plotted against the number of total recombinants within the same sliding window. A Spearman correlation coefficient was calculated between the recombinant count and the MFED values on either strand.

Taken together, these studies suggest that recombination does not preferentially occur within regions of RNA structure or sequence identity between templates. We propose that recombination *per se* is a random process, but is followed by a rigorous selection for ‘functional’ genomes [18, 26]. It is the analysis of the latter that has produced the prevailing view of the importance of sequence identity or structure in their generation, whereas in this study - specifically examining early recombinant genomes - we cannot find evidence supporting their involvement.

## Discussion

Recombination is an important evolutionary process in the single-stranded positive-sense RNA viruses [27, 28]. It is probably ubiquitous, and may be a necessary rescue mechanism to accommodate – and escape – the build-up of deleterious mutations that arise from the rapidly-processive [29], error-prone polymerase [30, 31]. Despite this, the molecular mechanisms underlying recombination remain relatively poorly understood. Several previous studies have implicated RNA secondary structure and particularly sequence similarity in determining the location of recombination junctions [12-14, 32, 33]. However, while it appears that this interpretation is widely accepted [10, 12, 23, 24], the studies reported here suggest that the early products of recombination are not significantly influenced by RNA sequence identity or structure, necessitating a rethink of the defining features of this important process.

By modifying our CRE-REP *in vitro* recombination assay [18], we investigated the influence of sequence identity or RNA structure in the parental genomes on the location of recombination junctions. We observed no significant variation in the distribution of recombinants occurring between the control region (Cluster 1) and the modified region (Cluster 2; **Fig 2**). Had either RNA sequence identity or structure been a significant influence on the mechanism of recombination we would have expected the distribution of recombinants to be skewed towards the Cluster region that favoured polymerase template-switching events. As all assays maintained a distribution similar to the wild type CRE-REP assay we inferred that neither sequence identity nor structure were a major influence on recombination.

By design, the CRE-REP assay generates viable progeny viruses for analysis and, as such, does not allow the initial population of hybrid genomes, from which viable recombinants are selected, to be characterised. We therefore went on to analyse the RNA population from HeLa cells co-infected with wild type PV1 and PV3. Despite the analysis of several hundred recombinant junctions by NGS (**Fig 3**) we could not detect any correlation between sequence identity (**Figs 4 and 5**) or RNA structure (**Fig 6**) and the enrichment of recombination junctions at any one particular site. Consequently, we propose that such regions undergo recombination at a rate no higher than would be expected by chance.

We have recently shown that the resolution of imprecise recombinants to wild type length genomes occurs via repeated template-switching events and is highly dependent on virus fitness [26]. This new understanding of resolution may help explain the discrepancy between the current prevailing view on recombination and the conclusions of this study if, as we suspect, the roles of sequence identity and structure are involved in genome function and fitness, rather than recombination *per se*. For example, studies that map crossover junctions to structured regions of recombinant genomes may conclude that they are causative, when the association may actually be correlative, and reflective of the genome modularity and relatively high fitness of a crossover in that specific region. Similarly, with genomes that exhibit variably extensive regions of sequence identity within an area in which recombinants are mapped, the functional selection may result in a clustering of crossovers in a region of sequence identity, not because of the identity *per se*, but because the resulting genome has enhanced fitness and is capable of competing better with others in the population.

A number of interesting questions remain about this important evolutionary mechanism which will require further study. If recombinants are generated at random as we propose, what are the predominant functional criteria that determine their selection? At a more mechanistic level, how does the template-switching event occur? Although our analysis strongly suggests that the process is random, there remains the possibility that regions of micro-identity may influence the process. There remains the interesting possibility that precise and imprecise junctions could be generated by independent mechanisms. We consider this unlikely; a sequence- and RNA structure-independent process - coupled with functional selection - can account for all of the range of crossover junctions we identify, all that have been reported in the literature, and would fit with the laws of parsimony.

## MATERIAL AND METHODS

### Virus and cell culture

HeLa cells and L929 mouse fibroblast cells were maintained in Dulbecco’s Modified Eagle Medium (DMEM) supplemented with 10% heat-inactivated FBS (FBS-DMEM). Poliovirus type 1 (Mahoney) and type 3 (Leon) were recovered following transfection of *in vitro* transcribed RNAs from full-length cDNAs. Additional virus stocks for replication competent PV3-2A/2B junction modified viruses FLC/PV3^L^, FLC/PV3^H^, and FLC/PV3^1^ (see below) were similarly generated and all virus stocks quantified by plaque assay on HeLa cells. Growth kinetics of viruses were determined following synchronous infection of HeLa cells at a multiplicity of infection (MOI) of 5 in serum-free media (SF-DMEM). Unabsorbed virus was removed by washing with PBS and cells incubated in fresh FBS-DMEM at 37°C, 5% CO2. Supernatants containing virus were harvested at various time points post-infection and quantified by plaque assay. Recombinant viruses isolated from CRE-REP assays were biologically cloned by limit dilution in 96-well plates of confluent HeLa cells. Virus-containing supernatant was removed after 3 days incubation at 37°C, 5% CO2 and stored at -80°C, and the remaining cell monolayer stained with a 0.1% crystal violet solution. Virus supernatants from the highest dilutions causing complete cytopathic effect (CPE) were utilised in further analysis

For co-infection studies, poliovirus type 1 and type 3 were used to co-infect confluent HeLa cells in four T175 flasks at an MOI of 10 of each virus. The cells were incubated for 30 minutes at 37°C before the virus was removed and the cells were washed with PBS. Fresh DMEM media was added and the flasks were incubated at 37°C for 5 h. After media removal, cells were washed with PBS, trypsinised and pooled in 2 ml DMEM media, giving a total of 7.6×10^7^ cells. The cells was lysed by three freeze-thaw cycles to extract virus and the debris was discarded. The supernatant contained the viruses was filtered through 0.2 μm filters and the resulting filtrate used for RNA extraction, followed by RT-PCR and NGS sequencing

### Design of modified 2A/2B junction sequences

A 450 nt sequence, equivalent to nucleotides 3599 to 4045, of both PV1 and PV3 was re-designed using the software package SSE [34] to generate sequences with the minimum and maximum amount of RNA structure possible by maintaining amino acid sequence and divergence between PV1 and PV3. The native PV1 and PV3 sequences were scrambled using the CDLR algorithm in SSE (**Fig S5**). This randomly scrambles the codon order, so disrupting any sequence-dependent underlying RNA structure, whilst maintaining the coding, codon usage and dinucleotide frequencies identical to those of the input sequence. CDLR randomisation was used to generate a large panel of variant sequences that retained these three characteristics. Each was processed using custom-written perl scripts to determine the sequence divergence from the input. Of those that were within 1% divergence (6230 PV1- and 6303 PV3-derived CDLR scrambles) the predicted RNA structure stability was calculated by determining the MFED (the minimal free energy difference [25]. MFED is a measure of the intrinsic folding energy of an RNA sequence and is determined by comparing an input sequence with 99 sequence-order randomised variants. The sequence identity of the 10 highest and lowest energy sequences was analysed and representative sequences with similar sequence identity were selected for construction as cDNAs (**Table S1**).

The same 450 nt sequence was altered for sequence identity by replacing the PV1 sequence with that of PV3 and *vice versa*. The sequence exchange was not 100% as there are 7 amino acid differences between PV1 and PV3 in this region – 2A residues 87, 116, 123, and 2B residues 3, 22, 26 and 30. At these amino acid positions the sequence was not altered between PV1 and PV3. Synthesis of all modified 450 nt DNA fragments was carried out by GeneArt (Life Technologies) and provided as sequence verified, plasmid DNA.

### CRE-REP assay, plasmids, in vitro RNA transcription and transfection

The CRE-REP assay used to generate recombinant viruses has been described previously [18]. Wild type cDNAs of PV1 donor template, pRLucWT, and PV3 acceptor template, pT7/SL3, were used as a background for the generation of the 2A/2B modified cDNAs. Standard molecular biology techniques were used to replace the 450 nt sequence spanning the 2A/2B boundary with the synthesised DNAs described above to generate the CRE-REP cDNA variants referred to as PV1^L^, PV1^H^, PV1^3^, PV3^L^, PV3^H^ and PV3^1^.

In addition, the relevant PV3 sequences were placed into a background of the replication competent PV3 cDNA, pT7/FLC, for growth kinetics analysis. All cDNAs were confirmed by sequence analysis prior to use. Plasmids were linearized with *Apa*I (pRLucWT background) or *Sal*I (pT7/SL3 or pT7/FLC background) and RNA transcribed using a HiScribe T7 High Yield RNA Synthesis kit (NEB) following the manufacturers’ protocol. RNA transcripts were DNaseI (NEB) treated to remove residual template DNA and column purified using a GeneJET RNA Purification Kit (ThermoFisher) prior to spectrophotometric quantification. For all CRE-REP variant assays equimolar amounts of both template RNAs (based on 250 ng of acceptor) were prepared with Lipofectamine 2000 (Life Technologies) in a 3:1 Lipofectamine 2000:RNA ratio as per manufacturers’ protocol, and transfected into 80% confluent L929 cell monolayers. Virus supernatant was recovered at 30 h post-transfection and virus quantified by plaque assay on HeLa cells. For virus stocks, 1 µg of RNA was transfected into 80% confluent HeLa cell monolayers using Lipofectamine 2000 as above, virus supernatant harvested at 12 h post-transfection and quantified by plaque assay on HeLa cells.

### RNA extraction and RT-PCR

Viral RNA was isolated from CRE-REP clarified cell culture supernatant samples using a GeneJET RNA Purification Kit (ThermoFisher) and reverse transcribed at 42°C using an oligo dT primer and SuperScript II reverse transcriptase (Life Technologies) as per manufacturers’ protocol. The region of recombination (VP1 to 2C) was amplified using primers PV3-F (5′-GCAAACATCTTCCAACCCGTCC-3′) and PV1-R (5′-TTGCTCTTGAACTGTATGTAGTTG-3′) and Taq polymerase (Life Technologies) with an initial denaturing at 94°C for 3 min, followed by 35 cycles of 94°C for 45 sec, 50°C for 30 sec and 72°C for 60 sec, and a final extension at 72°C for 10 min. Following poliovirus co-infection, total RNA was extracted from cells by QIAamp Viral RNA Mini Kit (Qiagen) and reverse transcribed as above. Subsequently 5 µl from the cDNA synthesis reaction was used in the PCR amplification. The region of recombination was amplified using primers PV3-F2 (5′-CTCCAAAGTCCGCATTTACA-3′) and PV1-R2 (5′-ATCAGGTTGGTTGCTACA-3′) and Taq polymerase (Promega) with an initial denaturing at 95°C for 2 min, followed by 35 cycles of 95°C for 1 min, 58.1°C for 30 sec and 72°C for 1.5 min, and a final extension at 72°C for 5 min.

### NGS library preparation

Illumina Nextera XT DNA kit was used to prepare the NGS samples for sequencing as per the manufacturer instructions. Briefly 1 ng from the amplicons was simultaneously fragmented and tagged with unique adapter sequences, followed by 10 cycles of PCR, before loading into the MiSeq instrument. The generated reads were uploaded to Illumina BaseSpace, a cloud-based genomics analysis and storage platform that directly integrates with all Illumina sequencers.

### Nomenclature

The terms precise and imprecise refer to whether the resulting recombinant genome is full length (precise) or contains insertions or deletions (imprecise) (**Fig S6**). We additionally defined the recombination junction as unambiguous, where the sequence between the donor and acceptor strands was dissimilar, or ambiguous where it was located within a short region of sequence identity in the parental genomes (**Fig S6**). We deliberately avoid use of the inexact term ‘homology’ which is variously used to indicate identity or relatedness. In two divergent recombining sequences, junctions can occur in regions of sequence identity or at positions where there is sufficient sequence divergence (whilst still being homologous) to unambiguously define the junction between the donor and acceptor sequences. Similarly, insertions or deletions can generate junctions where the origin of the sequences around the junction cannot be definitively assigned to the donor or recipient strand due to limited sequence identity around the junction.

## DATA AVAILABILITY

The raw sequencing data is available in the NCBI sequence read archive (SRA) with accession number: PRJNA648771

## SUPPLEMENTARY FIGURES

**Fig S1.**
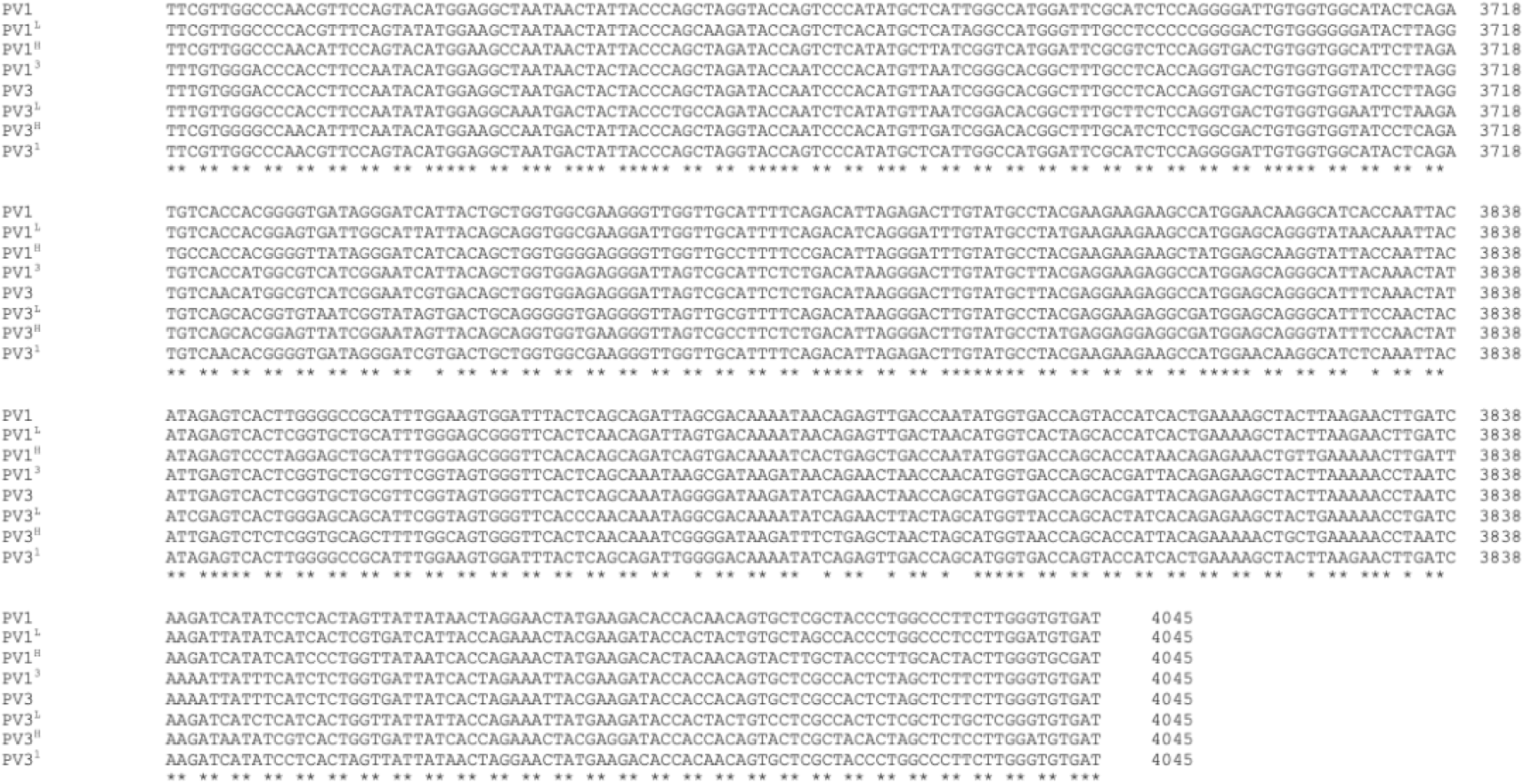
Sequence alignments of modified templates. The modified Cluster 2 sequences were aligned against parental PV1 and PV3 sequences using Clustal Omega.

**Fig S2.**
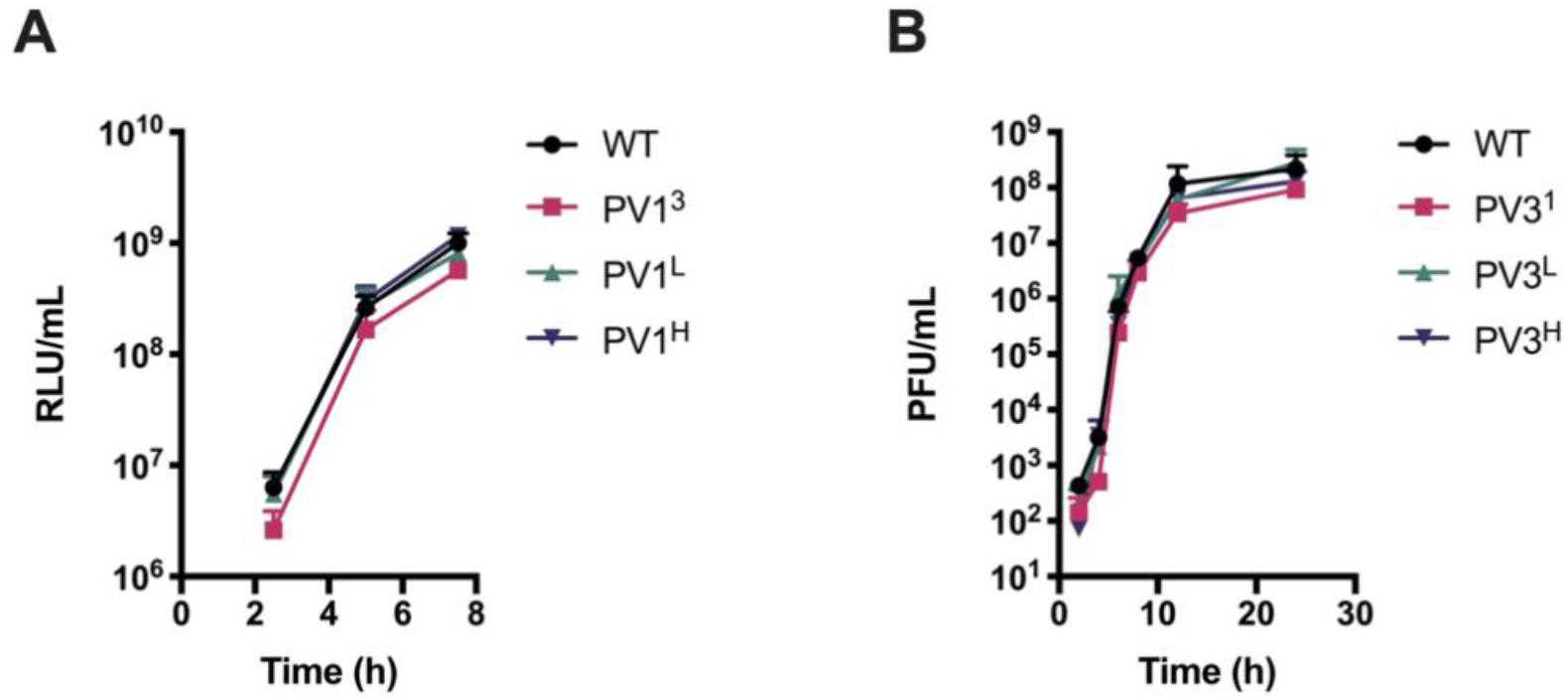
CRE-REP modified genome showed comparable characteristics to WT CRE-REP geneoms. (A) Replication kinetics for donor RNAs. HeLa cells were transfected with 200 ng RNA/well in 24-well plates and luciferase activity measured over an 8 h time course. Error bars represent the standard deviation of three experiments with samples assayed in triplicate. (B) Replication kinetics for acceptor RNAs. HeLa cells were infected with virus stock at MOI of 5 and virus-containing supernatants harvested over 24 h. Virus titres were determined by plaque assay on HeLa cells. Error bars represent the standard deviation of three experiments.

**Fig S3.**
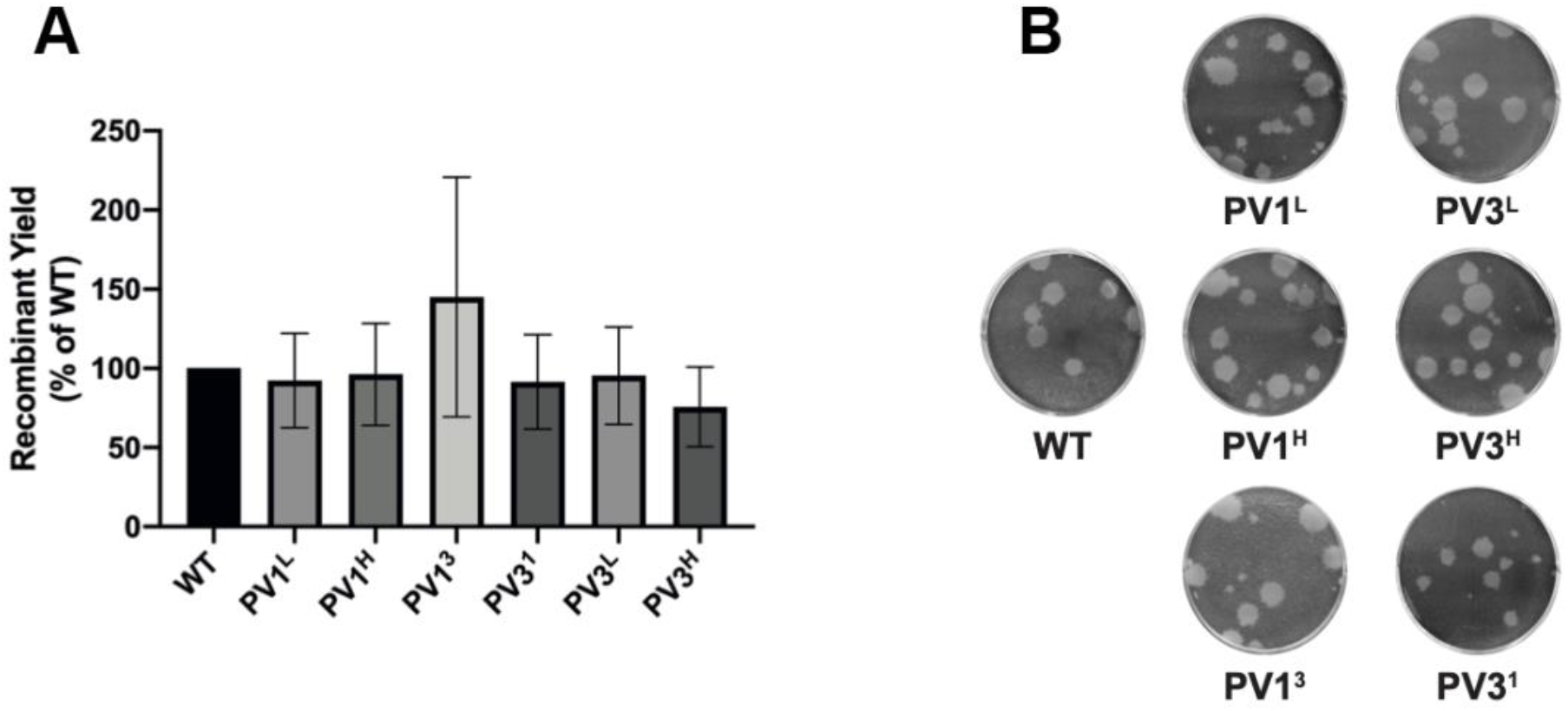
Quantification and characterisation of donor and acceptor RNAs in modified CRE-REP assays. (A) Recombinant yield from modified CRE-REP assays. RNAs were co-transfected into L929 cells in equimolar amounts and virus-containing supernatants were harvested 30 h post-transfection. Recombinant virus yields were determined by plaque assay on HeLa cells and expressed as a percentage of the WT assay. Error bars represent standard error of the mean of three co-transfection experiments. All modified assays were not significantly different to WT, as determined by unpaired t-test. (B) Plaque morphology comparison between recombinant viruses obtained from each of the CRE-REP assays.

**Fig S4.**
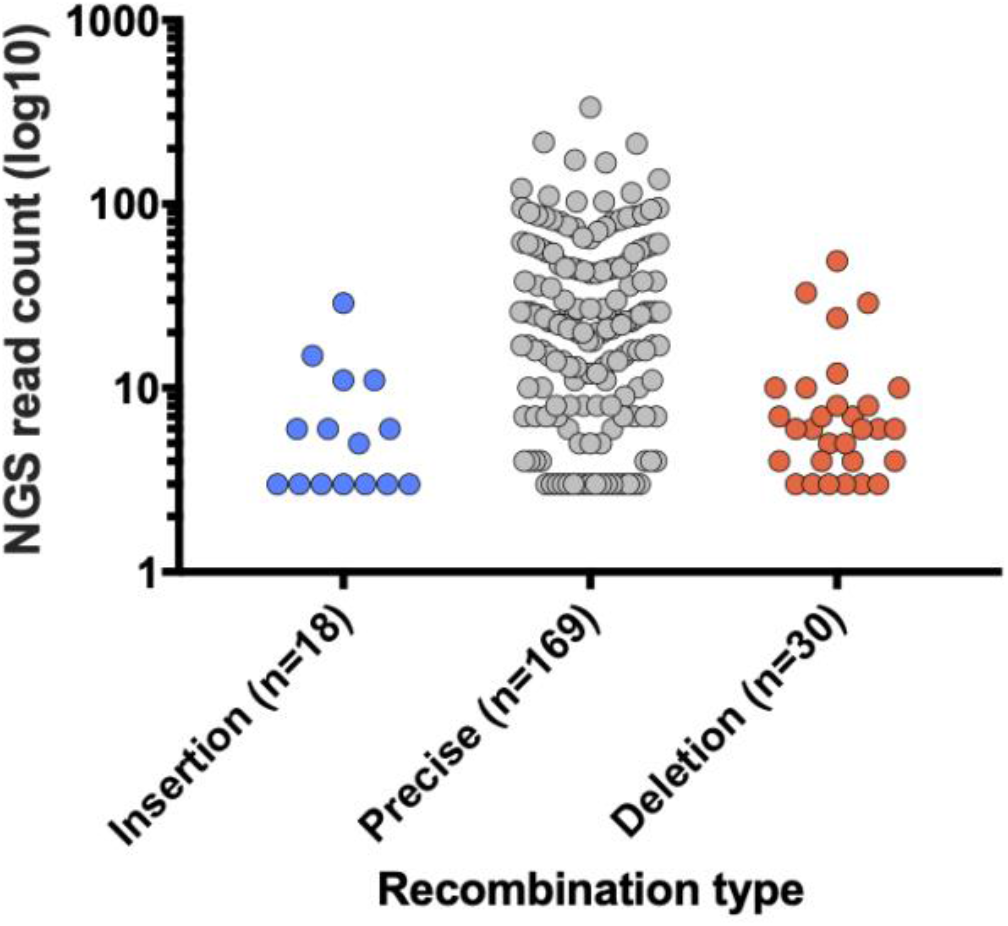
Sequencing analysis of the co-infection technical replicate sample. The number of precise and imprecise (insertion and deletion) recombinants was plotted against the number of NGS reads (left panel). Each dot represents a unique recombinant.

**Fig S5.**
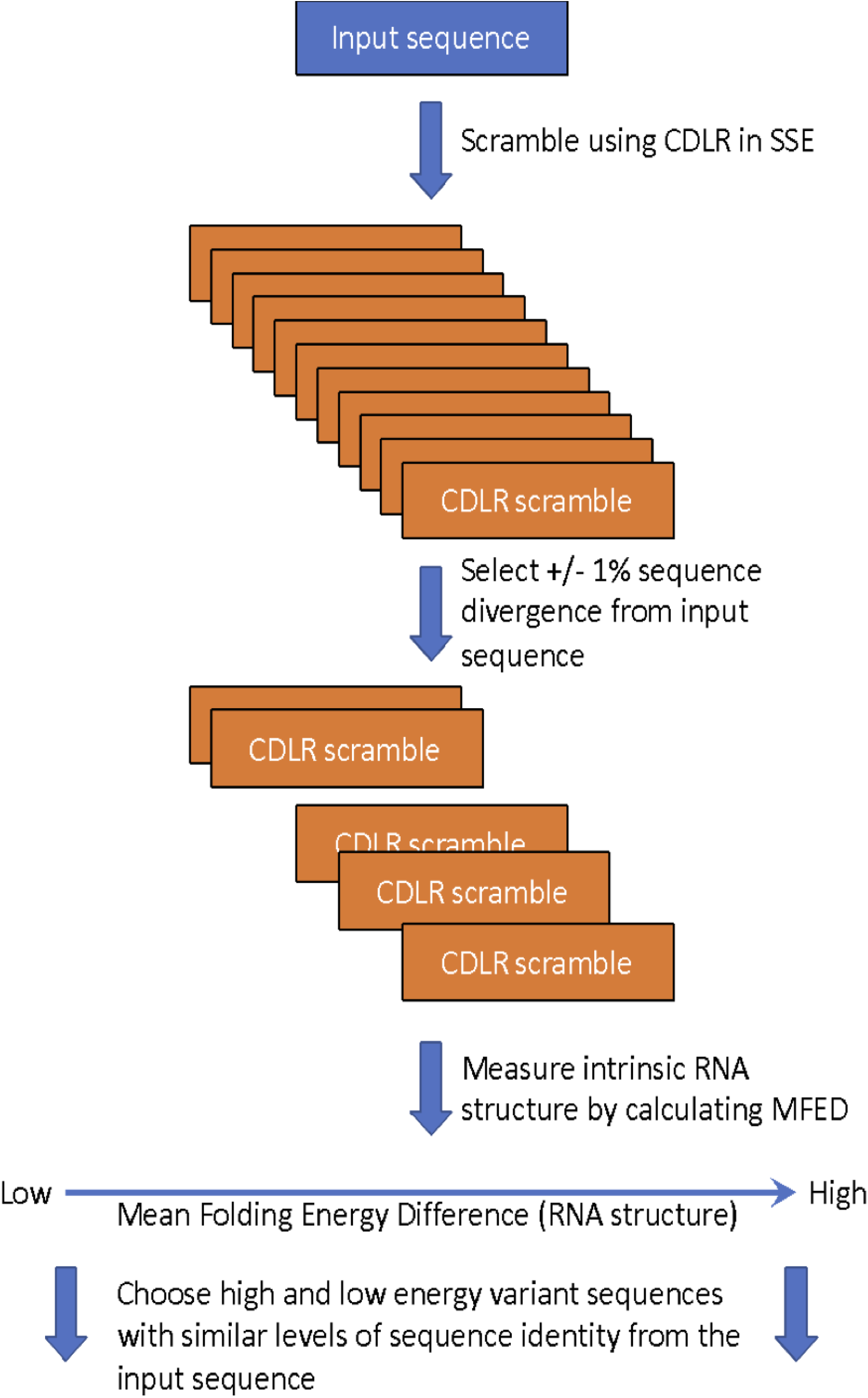
Design flowchart for RNA sequences with high or low levels of RNA structure. The input sequences were scrambled using the CDLR algorithm within the SSE package (reference 25). This method scrambles codon order, but maintains the open reading frame, the encoded polyprotein, the dinucleotide and codon frequencies. Scrambled sequences within +/-1% sequence divergence from the input sequence were selected. Each was analysed and ordered for inherent RNA structure by calculating the mean folding energy difference (MFED) between the CDLR scrambled sequences and 99 sequence-order randomised variants (see reference 34). Representative high and low structured variants, both exhibiting similar sequence identity to the native input poliovirus sequence, were selected from the 10 sequences with the highest or lowest level of intrinsic structure respectively.

**Fig S6.**
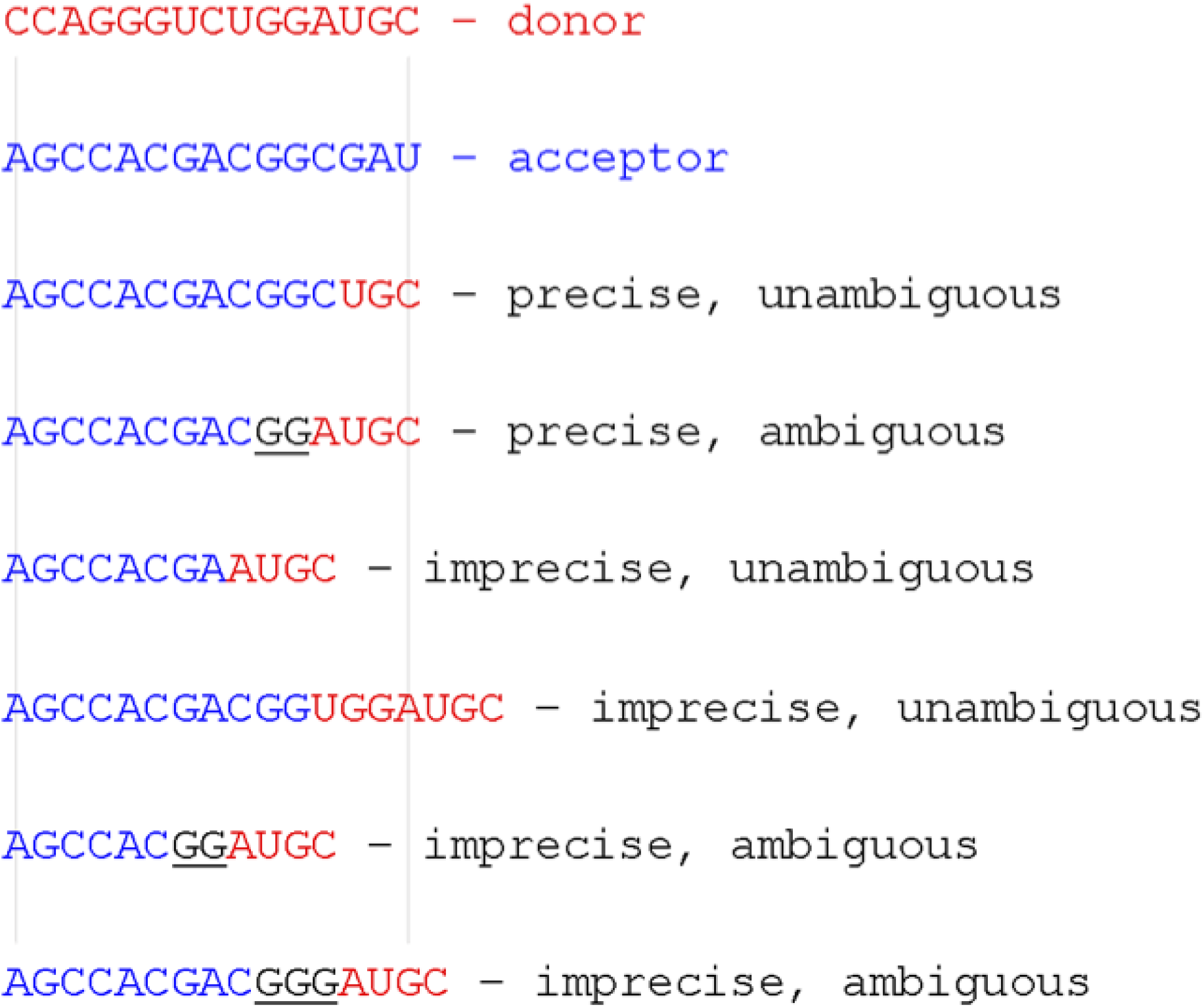
Recombination junction definition. Donor and recipient sequences are shown in red and blue respectively. Precise junctions have no insertions or deletions with respect to the length of the parental sequences. Imprecise junctions contain insertions or have sequences deleted. The position of unambiguous junctions can be defined by the position of the join in the donor and acceptor sequences. Ambiguous junctions, whether in precise or imprecise junctions, cannot be exactly defined due to limited sequence identity between donor and acceptor at the junction, and are highlighted in <u>black underlined</u> text. The two thin vertical lines refer to start and end of the precise recombinants, to visually aid reading the figure.

**Table S1.** Percentage sequence identity of structure modified sequences.

|  | PV1 | PV1<sup>L</sup> | PV1<sup>H</sup> | PV3 | PV3<sup>L</sup> | PV3<sup>H</sup> |
| --- | --- | --- | --- | --- | --- | --- |
| PV1 | - | 85.46% | 85.46% | 77.63% | 78.52% | 77.40% |
| PV3 | 77.63% | 78.75% | 76.96% | - | 85.46% | 85.46% |
| $\Delta G$<br>(kcal/mol) | -124.90 | -112.30 | -149.90 | -140.00 | -126.20 | -149.80 |

**Table S2.**
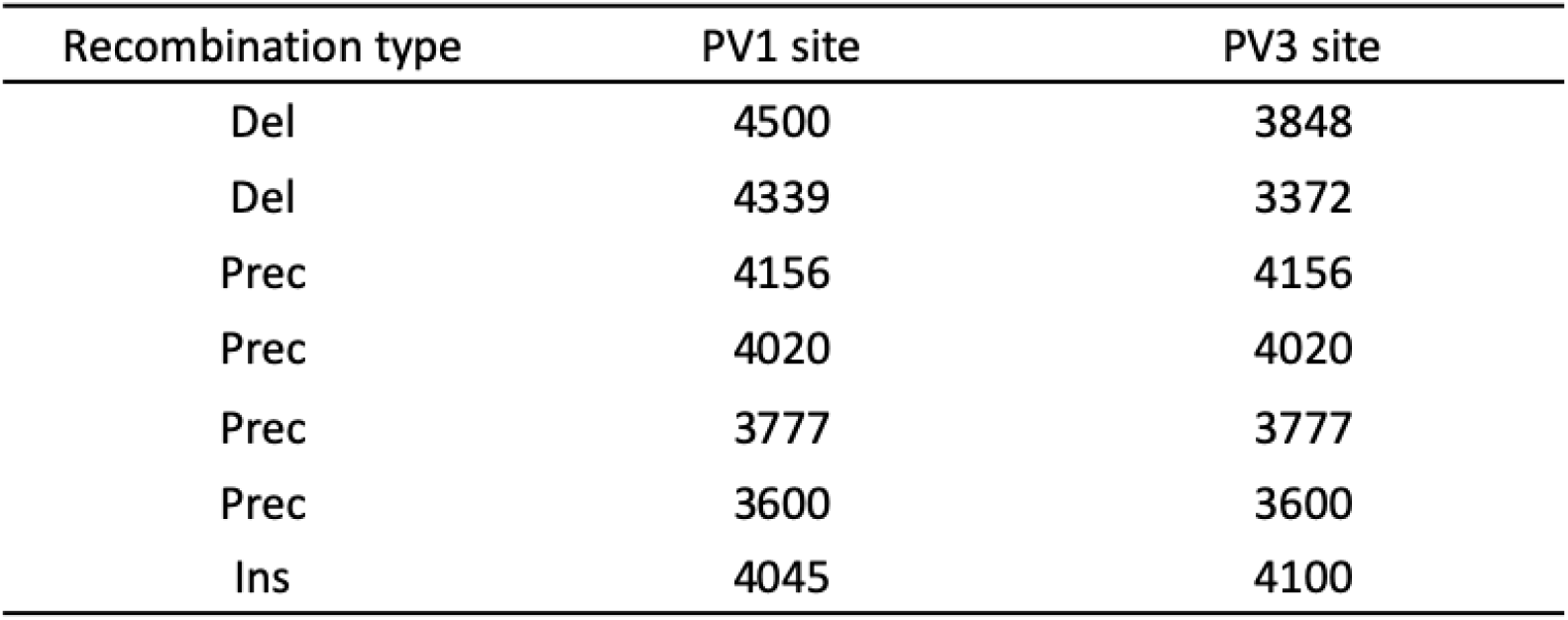
Recombination junctions isolated by PCR cloning and sequenced by Sanger-sequencing. (Del = Deletion, Prec = Precise, Ins = Insertion)

## Reference

1. Hahn, C.S., et al., Western equine encephalitis virus is a recombinant virus. Proc Natl Acad Sci U S A, 1988. 85(16): p. 5997–6001.

2. Ge, X.Y., et al., Isolation and characterization of a bat SARS-like coronavirus that uses the ACE2 receptor. Nature, 2013. 503(7477): p. 535–8.

3. Li, K.S., et al., Genesis of a highly pathogenic and potentially pandemic H5N1 influenza virus in eastern Asia. Nature, 2004. 430(6996): p. 209–13.

4. Smith, G.J., et al., Origins and evolutionary genomics of the 2009 swine-origin H1N1 influenza A epidemic. Nature, 2009. 459(7250): p. 1122–5.

5. Gmyl, A.P., et al., Nonreplicative RNA recombination in poliovirus. J Virol, 1999. 73(11): p. 8958–65.

6. Gmyl, A.P., et al., Nonreplicative homologous RNA recombination: promiscuous joining of RNA pieces? Rna, 2003. 9(10): p. 1221–31.

7. King, A.M., Preferred sites of recombination in poliovirus RNA: an analysis of 40 intertypic cross-over sequences. Nucleic Acids Res, 1988. 16(24): p. 11705–23.

8. Kirkegaard, K. and D. Baltimore, The mechanism of RNA recombination in poliovirus. Cell, 1986. 47(3): p. 433–43.

9. Woodman, A., et al., Biochemical and genetic analysis of the role of the viral polymerase in enterovirus recombination. Nucleic Acids Res, 2016. 44(14): p. 6883–95.

10. Nagy, P.D. and J.J. Bujarski, Efficient system of homologous RNA recombination in brome mosaic virus: sequence and structure requirements and accuracy of crossovers. J Virol, 1995. 69(1): p. 131–40.

11. Sergiescu, D., A. Aubert-Combiescu, and R. Crainic, Recombination between guanidine-resistant and dextran sulfate-resistant mutants of type 1 poliovirus. J Virol, 1969. 3(3): p. 326–30.

12. White, K.A. and T.J. Morris, RNA determinants of junction site selection in RNA virus recombinants and defective interfering RNAs. RNA, 1995. 1(10): p. 1029–40.

13. Nagy, P.D., C. Zhang, and A.E. Simon, Dissecting RNA recombination in vitro: role of RNA sequences and the viral replicase. EMBO J, 1998. 17(8): p. 2392–403.

14. Carpenter, C.D., et al., Involvement of a stem-loop structure in the location of junction sites in viral RNA recombination. J Mol Biol, 1995. 245(5): p. 608–22.

15. Bruyere, A., et al., Frequent homologous recombination events between molecules of one RNA component in a multipartite RNA virus. J Virol, 2000. 74(9): p. 4214–9.

16. Bujarski, J.J. and P. Kaesberg, Genetic recombination between RNA components of a multipartite plant virus. Nature, 1986. 321(6069): p. 528–31.

17. Rowe, C.L., et al., Generation of coronavirus spike deletion variants by high-frequency recombination at regions of predicted RNA secondary structure. J Virol, 1997. 71(8): p. 6183–90.

18. Lowry, K., et al., Recombination in enteroviruses is a biphasic replicative process involving the generation of greater-than genome length ‘imprecise’ intermediates. PLoS Pathog, 2014. 10(6): p. e1004191.

19. Egger, D. and K. Bienz, Intracellular location and translocation of silent and active poliovirus replication complexes. J Gen Virol, 2005. 86(Pt 3): p. 707–18.

20. Bowtie 2: Manual. 2016; Available from: http://bowtie-bio.sourceforge.net/bowtie2/manual.shtml#bowtie2-options-score-min.

21. Routh, A. and J.E. Johnson, Discovery of functional genomic motifs in viruses with ViReMa-a Virus Recombination Mapper-for analysis of next-generation sequencing data. Nucleic Acids Res, 2014. 42(2): p. e11.

22. Alnaji, F.G., et al., Sequencing Framework for the Sensitive Detection and Precise Mapping of Defective Interfering Particle-Associated Deletions across Influenza A and B Viruses. J Virol, 2019. 93(11).

23. Figlerowicz, M., Role of RNA structure in non-homologous recombination between genomic molecules of brome mosaic virus. Nucleic Acids Res, 2000. 28(8): p. 1714–23.

24. Cascone, P.J., T.F. Haydar, and A.E. Simon, Sequences and structures required for recombination between virus-associated RNAs. Science, 1993. 260(5109): p. 801–5.

25. Simmonds, P., A. Tuplin, and D.J. Evans, Detection of genome-scale ordered RNA structure (GORS) in genomes of positive-stranded RNA viruses: Implications for virus evolution and host persistence. RNA, 2004. 10(9): p. 1337–51.

26. Bentley, K., et al., Imprecise recombinant viruses evolve via a fitness-driven, iterative process of polymerase template-switching events. PLoS Pathog, 2021. 17(8): p. e1009676.

27. Simon-Loriere, E. and E.C. Holmes, Why do RNA viruses recombine? Nat Rev Microbiol, 2011. 9(8): p. 617–26.

28. Worobey, M. and E.C. Holmes, Evolutionary aspects of recombination in RNA viruses. J Gen Virol, 1999. 80 (Pt 10): p. 2535–43.

29. Duffy, S., Why are RNA virus mutation rates so damn high? PLoS Biol, 2018. 16(8): p. e3000003.

30. Kempf, B.J., et al., Picornavirus RNA Recombination Counteracts Error Catastrophe. J Virol, 2019. 93(14).

31. Crotty, S., C.E. Cameron, and R. Andino, RNA virus error catastrophe: direct molecular test by using ribavirin. Proc Natl Acad Sci U S A, 2001. 98(12): p. 6895–900.

32. Bentley, K. and D.J. Evans, Mechanisms and consequences of positive-strand RNA virus recombination. J Gen Virol, 2018. 99(10): p. 1345–1356.

33. Muslin, C., et al., Recombination in Enteroviruses, a Multi-Step Modular Evolutionary Process. Viruses, 2019. 11(9).

34. Simmonds, P., SSE: a nucleotide and amino acid sequence analysis platform. BMC Res Notes, 2012. 5: p. 50.

